# Mesaconic acid delivery via biomaterials and engineered probiotics to the gut reduces inflammation in Crohn’s Diseases mouse models

**DOI:** 10.64898/2026.10.01.756006

**Authors:** Huikang Qian, Drishtant Singh, Zixuan Cheng, Wenfei Zhao, Bo Zhang, Taravat Khodaei, Abhirami Thumsi, Sanmoy Pathak, Abhirami P. Suresh, Paola Menghini, Julia Schiek, Viveka Rabara, Carlo De Salvo, Harihara Baskaran, Yuji Aso, Madhan Mohan Chandra Sekhar Jaggarapu, Abigail Raffner Basson, Abhinav P. Acharya

## Abstract

Mesaconate (MES, an immunosuppressive metabolite) but not itaconate is upregulated in stool of Crohn’s Disease (CD) patients, albeit below therapeutic levels.. However, colonic MES level is difficult to maintain because of its high absorption rate, limiting its therapeutic potential for CD. Herein, we developed two strategies that elevate MES level and suppress inflammation: EcN-CAD, an engineered probiotic Escherichia coli Nissle 1917 (EcN) that overexpresses cis-aconitate decarboxylase (CAD, produces itaconic acid that is converted to MES); and MES+Zn, a Zinc-MES ionically crosslinked formulation. Mechanistically, MES modulates T cell function by downregulating glycolytic and mitochondrial activity and decrease RORγt expression. In preclinical SAMP1/YitFC dexamethasone-withdrawal model and TNFDARE model, MES+Zn decreased gut permeability significantly compared to controls and reduced myeloperoxidase (MPO, an inflammatory marker) 1000-fold compared to controls. EcN-CAD probiotics decreased RORγt expression in CD4^+^ and CD8^+^ T cells, gut permeability and inflammation in preventive SAMP1/YitFC model, and SAMP1/YitFC dexamethasone-withdrawal model. Overall, MES rewires T cell responses and technologies that enhance MES levels can be effective in gut-related inflammatory diseases.

## Main

The inflammatory bowel disease (IBD) subtype Crohn’s disease (CD) is a chronic, relapsing inflammatory condition of the gastrointestinal tract that affects an estimated 1 million individuals in the United States^1,2^, with both its incidence and prevalence continuing to rise^1,2^. CD is characterized by discontinuous transmural intestinal inflammation, impaired gut barrier function (i.e. leaky gut syndrome), and a high cumulative risk of surgery, with up to 70% of patients requiring surgical intervention within 10 years of diagnosis^2^. Despite advances in biologic therapies targeting TNF-α, IL-12/23, and integrins, many patients experience primary non-response or secondary loss of response over time ^2,3^. Treatments are further limited by high cost and significant side effects including increased infection risk, malignancy and other adverse effects arising from systemic immunosuppression ^4,5^.

In CD, dysregulated T-cell responses are central drivers of persistent intestinal inflammation, particularly involving an imbalance favouring pro-inflammatory CD4 T helper subsets (T_h_1, T_h_17)^6,7^. Enhanced differentiation of naïve CD4 T cells into T_h_17 cells is characterised by strong upregulation of retinoic acid receptor-related orphan nuclear receptor γt (RORγt), which directly drives expression of the pro-inflammatory cytokines IL-17A, IL-17F, and IL-22. Collectively, this promotes mucosal inflammation, epithelial barrier disruption, and recruitment of inflammatory leukocytes into intestinal tissues^6,8–12^. CD is also characterized by decreased Treg cell frequency, which is responsible for suppression of pathogenic effector T-cell responses^11–15^. Additionally, activated CD8 cytotoxic T cells accumulate within inflamed intestinal mucosa, exacerbating tissue damage through cytotoxic mechanisms involving perforin, granzyme B, and pro-inflammatory cytokines (IFN-γ, TNF-α), as well as perpetuating inflammation via tissue-resident memory populations^11,16^.

Use of immune-modulating metabolites to reprogram immune cells has been widely studied for their therapeutic potential in CD management^17–19^. Itaconic acid (ITA), a metabolite produced by immune-responsive gene 1 (ACOD1/IRG1, also known as CAD) in activated macrophages, has reported immunoregulatory and antioxidative properties in IBD, including CD^18–23^. ITA is generated by cis-aconitate decarboxylase from cis-aconitate, an intermediate metabolite in tricarboxylic acid cycle. In T cells, ITA inhibits glycolysis and oxidative phosphorylation by decreasing the intracellular levels of S-adenosylmethionine (SAM) and 2-hydroxyglutarate (2-HG)^20^. The decreased SAM/SAH ratio in T_h_17 cells lead to hypomethylation at the IL17a promoter site, suppressing IL-17A expression, while decreased 2-HG levels in T_reg_ cells promote demethylation at the Foxp3 locus, enhancing FOXP3 expression^21^.

Mesaconic acid (MES) and citraconic acid (CIT) are known isomers of ITA and exhibit similar immune suppressive effects on macrophages^21,22^. MES is endogenously synthesized from ITA during LPS activation, and similar to ITA, lowers glycolytic flux and suppresses pro-inflammatory cytokine release, such as IL-6, TNF-a and IL-12 in macrophages^21,22^. Unlike ITA, MES leaves succinate-dehydrogenase (SDH) activity and mitochondrial respiration largely untouched and exerts its effects independently of ATF3^21^. Whether MES exerts comparable effects in adaptive immune cells, and whether it contributes to intestinal immune regulation, is unknown. Herein we show that MES downregulates mitochondrial respiration and RORγt expression in T cells, supporting its therapeutic potential in CD. We also show that fecal levels of MES (but not ITA) correlate with disease severity in human CD patients. To address issues surrounding small metabolite absorption and clearance in the intestine we developed a genetically engineered *Escherichia coli Nissle* 1917 probiotic engineered to express cis-aconitate decarboxylase (CAD), yielding a strain that produces MES in situ. However, since soluble small metabolites are rapidly diluted, absorbed and cleared in the intestinal lumen, making sustained exposure at inflamed mucosal surfaces difficult to achieve we also developed zinc-mesaconic acid (MES+Zn) formulation to prolong persistence in the gut. Across patient stool measurements, primary T-cell assays, proteomics and multiple mouse models of ileitis, we show that MES positively correlate with inflammatory markers in CD, suppresses pro-inflammatory T-cells and reduces mitochondrial and glycolytic activity in activated T cells. *In vivo*, MES+Zn improved inflammatory and epithelial barrier readouts in a TNFa driven acute model and a spontaneous chronic CD remission model. Additionally, EcN-CAD also reduced permeability, fecal myeloperoxidase (MPO), pro-inflammatory T-cell markers, and tissue-injury features across treatment, remission-maintenance and prophylactic settings. Together, these data establish MES delivery as a tractable immunoengineering strategy for intestinal inflammation and identify EcN-CAD as a preclinical prototype for programmable gut-localized metabolite therapy.

## Results

### Fecal mesaconate levels positively correlate with fecal myeloperoxidase, a marker of intestinal inflammation in Crohn’s disease

To evaluate the clinical relevance of MES in CD, we quantified fecal MES levels from patients with CD and non-IBD controls using LC-MS/MS (**Fig. 1a-c**). In parallel, intestinal inflammation was assessed by measuring fecal myeloperoxidase (MPO), a validated marker of neutrophil infiltration, and mucosal inflammation^28^. Across CD patient samples, fecal MES levels increased with rising MPO levels, revealing a positive correlation between MES abundance and intestinal inflammatory burden with P value of 0.009 (Pearson’s correlation coefficient = 0.47) (**Fig. 1b**). In contrast, no significant correlation was observed between MPO and itaconate (ITA) (Pearson’s correlation coefficient = –0.05, P value of 0.79) (**Fig. 1c**). These data indicate that stool MES, but not ITA, is an indicator of inflammatory activity in the CD patients. The possible cause of this result might be that ITA released in gut was consumed by gut microbiota and catabolized into MES.

**Figure 1:**
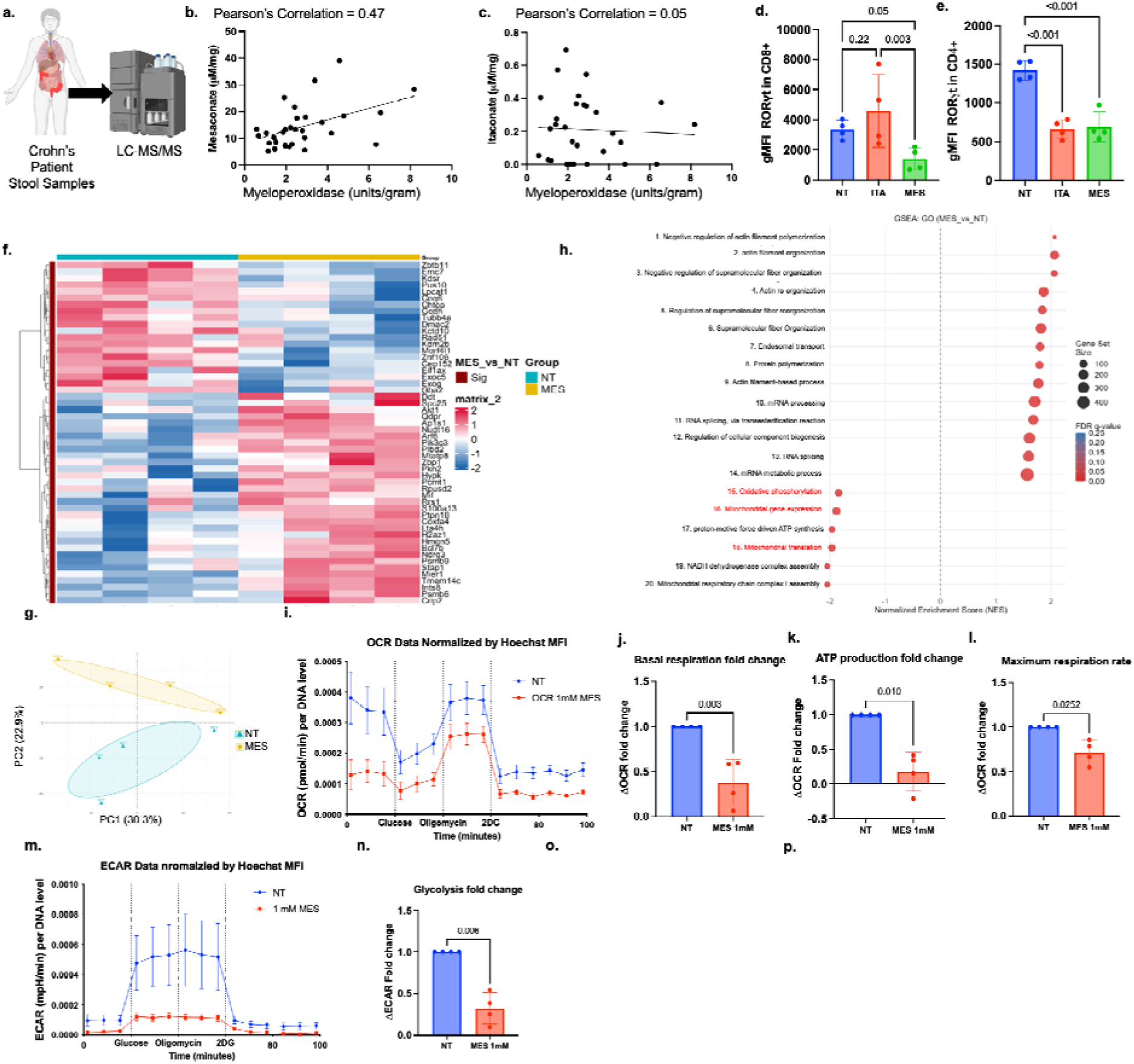
Fecal MES tracks intestinal inflammation in Crohn’s disease and MES suppresses mitochondrial metabolism and RORgt expression in T cells. (a) Quantification by LC-MS/MS of MES and ITA levels in CD patient fecal samples. (b-c) Pearson’s correlation between MES and ITA level and MPO in CD patients’ fecal samples. n = 30 patients. (d-e) gMFI of RORgt expression in CD8^+^ and CD4^+^ T cells when spleen CD3^+^ T cells were treated with 1mM MES. One-way ANOVA, Fisher LSD post-hoc analysis n = 4 mice. (f-h) Global proteomic of activated T cell treated with 1mM MES or untreated. n = 4 mice. (f) Heatmap shows the relative abundance of LCQ intensity of differentially expressed proteins in activated T cell treated with MES or untreated. (g) Principal-component analysis of activated T cells. (h) Gene set enrichment analysis (GSEA) of gene ontology (GO) pathway comparing between MES-treated and untreated T cells. (i-n) Seahorse extracellular flux assay of activated spleen CD3^+^ T cell treated with 1mM MES or untreated normalized by DNA level. Student’s paired two-tailed t-test, N = 4 mice.

### Effects of MES on T cell inflammatory markers *in vitro*

MES has been determined to have immunomodulatory effects on macrophages, however the effect on T cells remain uninvestigated ^21^. To understand how MES affects T cell responses, we introduced MES and ITA to activated CD3^+^ T cells. Notably, MES treatment resulted in a significant reduction in RORγt expression in CD8□T cells compared with untreated controls in a dose dependent manner (**Fig. 1d, S1a**). A similar reduction in RORγt expression was observed in CD4□T cells following MES treatment across multiple concentrations (**Fig. 1e, S1b**). In contrast, ITA produced effects on RORγt expression in CD8□ were not significantly different than controls, and ITA had a similar result as MES on CD4□T-cell populations. To understand if MES exerts anti-inflammatory effects on cytokine level, we treated CD3^+^ T cells isolated from spleens of C57BL/6J mice with MES and activated cells with anti-CD3 and anti-CD28. Interestingly, we observed reduced secretion of IL-17 (**Fig S1d**). To investigate the protein level change of MES-mediated suppression of inflammatory T-cell responses, we performed global proteomic analysis on activated mice T cells treated with MES or left untreated (NT). Heatmap (**Fig. 1f**) and Principal-Component Analysis (**Fig S1.c)** showed clear separation between MES-treated and untreated samples, indicating that MES induced a distinct proteomic state in activated T cells and differential abundance analysis further identified multiple proteins altered by MES treatment. To determine whether these changes converged on specific biological programs, we performed gene set enrichment analysis (GSEA) using Gene Ontology (GO) biological process terms. MES-treated cells showed positive enrichment of pathways related to negative regulation of actin filament polymerization, actin cytoskeleton organization, actin filament-based processes and supramolecular fiber organization, indicating that MES altered cytoskeletal programs associated with T-cell activation and structural remodeling (**Fig. 1h**). In parallel, MES-treated cells showed negative enrichment of oxidative phosphorylation (q value = 0.007), mitochondrial gene expression (q value = 0.005), proton motive force-driven ATP synthesis (q value = 0.007), mitochondrial translation (q value = 0.006), NADH dehydrogenase complex assembly (q value = 0.007) and mitochondrial respiratory chain complex I assembly (q value = 0.010), supporting the hypothesis that MES suppresses mitochondrial bioenergetic programs (**Fig. 1h**). Together, these data show that MES treatment is associated with coordinated remodeling of actin-associated pathways and attenuation of mitochondrial oxidative metabolism in activated T cells, providing a mechanistic framework for the reduced RORγt expression and IL-17 production observed after MES treatment. To determine whether MES directly alters T-cell metabolism, activated CD3□T cells were treated with 1 mM MES (concentration that showed decreased RORγt expression **Fig. 1d, S1a, b)** and analyzed using extracellular flux assays. Consistent with this shift, MES treatment reduced oxygen consumption rate (OCR) across the mitochondrial stress test compared with untreated controls (**Fig. 1i**). Quantification of OCR-derived parameters showed that MES significantly decreased basal respiration, ATP-linked respiration, and maximal respiration relative to untreated cells (**Fig. 1j-l**). These data suggest that MES limits mitochondrial respiratory activity in activated T cells. MES also strongly reduced the extracellular acidification rate (ECAR), indicating suppression of glycolytic metabolism (**Fig. 1m**). Following glucose and oligomycin injection, untreated activated T cells showed a robust increase in ECAR, whereas MES-treated cells exhibited a markedly blunted glycolytic response. Quantification confirmed that MES significantly reduced glycolysis compared with untreated controls (**Fig. 1n**). Together with the OCR data, these results indicate that MES does not simply shift activated T cells from oxidative phosphorylation toward glycolysis, but instead broadly suppresses both major energy-producing pathways. These findings suggest that MES metabolically restrains activated T cells and preferentially suppresses RORγt-associated inflammatory programming in CD8 T cells, supporting its role as an immunometabolic regulator in the context of intestinal inflammation.

### MES and MES+Zn decrease pro-inflammatory marker expression in TNFΔARE mice and in SAMP dexamethasone remission model

Many CD patients relapse after corticosteroid-induced remission. Thus, to better reflect the clinical scenario of post-steroid remission we used a dexamethasone (DEX) remission model^29^ (equivalent to corticosteroid-induced remission in CD patients) using two genetically distinct mouse models of CD-ileitis; TNFΔARE and SAMP mice. Our DEX remission model decreases ileitis severity by ∼70%. However, ileitis recurs as early as 2 weeks after DEX cessation, enabling evaluation of therapies that maintain, but do not induce, remission. Given that MES downregulates TNFa expression in macrophages^22^, we first used TNFDARE mice, a TNF-driven acute model of intestinal inflammation, to evaluate the ability of MES to prolong DEX-induced inflammation relief. Interestingly, we found that the salt formulation MES+Zn could prolong MES retention time in the gut compared to all other groups, and about 7 times higher in both TNFΔARE mice and SAMP mice stool samples when compared to MES alone group (**Fig. 2d, g**). These data highlight the importance of utilizing Zn as a complexing agent to enhance MES presence in the colon. Moreover, MES+Zn was characterised by FTIR and ¹H NMR (**Fig. S1f, g**). Compared to MES, MES+Zn showed upfield shifts of both the vinyl (δ 6.80 → 6.47 ppm) and methyl (δ 2.16 → 1.93 ppm) resonances, which demonstrates that MES is non-covalently bound to Zn. The FTIR spectrum of MES+Zn lacks the free carboxylic acid C=O band and dominated by paired asymmetric and symmetric carboxylate stretches, consistent with the fully deprotonated carboxylate form. Spectra from two batches synthesised two years apart were superimposable, indicating reproducible synthesis and stability of the formulation for at least 24 months at room temperature (21 °C).

**Figure 2:**
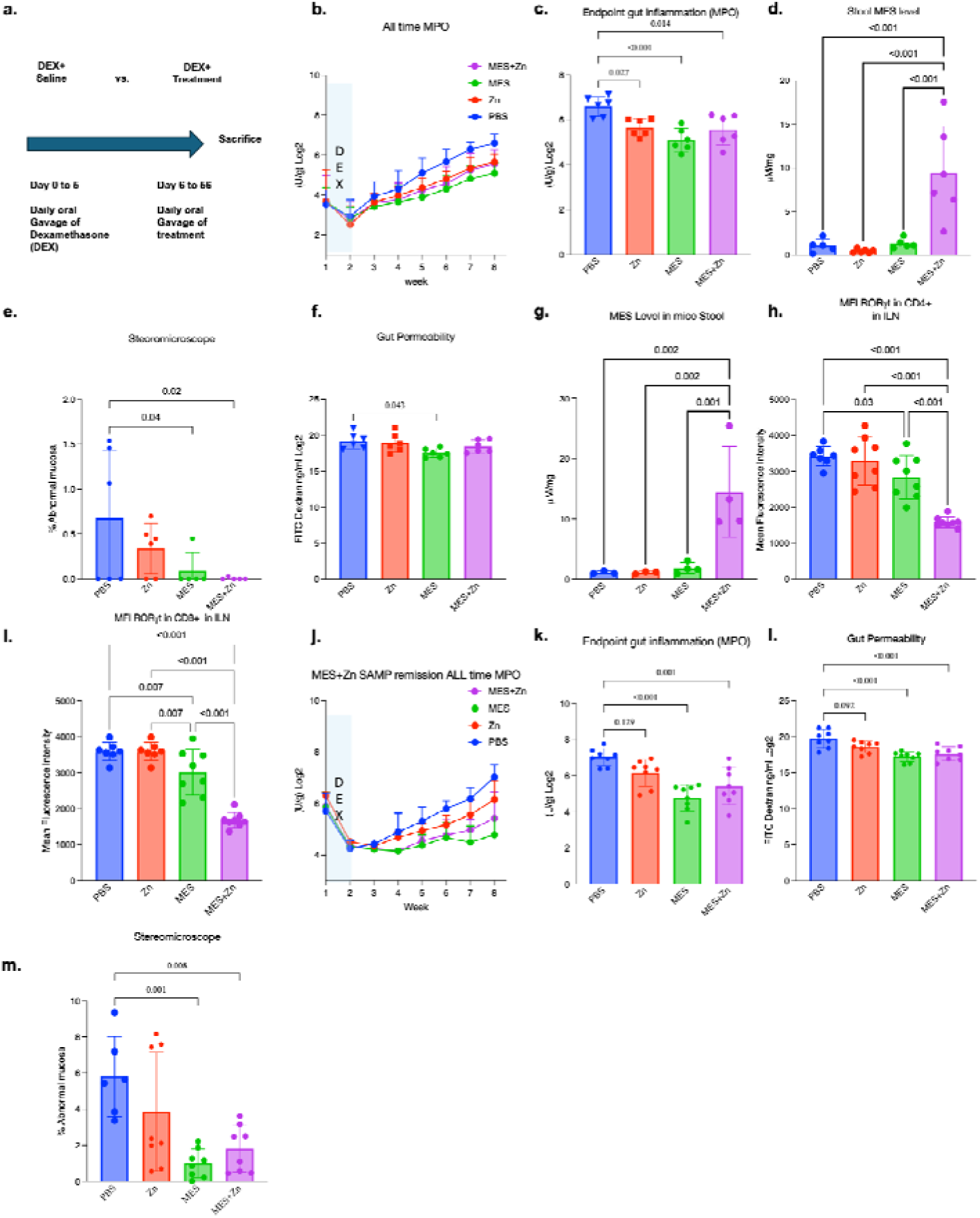
MES and MES+Zn protect TNFDARE and SAMP mice gut barrier and reduces RORγt expression in T cells. **(a)** TNFDARE & SAMP mice were given MES or MES+Zn orally for 50 days after 5 days of DEX treatment. **(b)** Weekly Fecal MPO in TNFDARE mice over 8 weeks **(c)** Fecal MPO at week 8 (sacrifice) in TNFDARE mice. **(d)** MES level in TNFDARE mice stool samples at sacrifice. **(e)** 3-D Stereomicroscopy of % abnormal mucosa at sacrifice in TNFDARE mice. **(f)** FITC-dextran gut permeability assay (*in vivo*). **(g)** MES level in SAMP mice stool samples at sacrifice. **(h-i)** gMFI of RORgt expression in mesenteric lymph node (MLN) CD4^+^ and CD8^+^ T cells. **(j)** Weekly Fecal MPO in SAMP mice over 8 weeks **(k)** Fecal MPO at week 8 (sacrifice) in SAMP mice. **(l)** FITC-dextran gut permeability assay at sacrifice. **(m)** 3-D Stereomicroscopy of % abnormal mucosa at sacrifice in SAMP mice. One-way ANOVA, Fisher’s LSD post-hoc test, n = 6 mice for TNFDARE and n = 8 mice for SPF-SAMP remission model.

To test if MES+Zn modulated symptoms in vivo, specific pathogen free (SPF) 14-week-old TNFΔARE (i.e. established ileitis) were administered with DEX to induce remission. Mice were then treated with PBS, Zn, MES, or MES+Zn for 50 days (**Fig. 2a**). Longitudinal measurements of fecal MPO showed that both MES and MES+Zn treatments slowed the increase in inflammatory burden over time compared with PBS-treated controls (**Fig. 2b**). At the study endpoint, fecal MPO levels were significantly reduced in both treatment groups (**Fig. 2c**). Consistent with these biochemical findings, 3D stereomicroscopy (3D-SM) scoring of % abnormal mucosa^28^ revealed reduced intestinal pathology in MES– and MES+Zn-treated mice significantly when compared to PBS controls (**Fig. 2e**). However, no significant difference was observed in histological scoring (**Fig. S4a, b)**. Assessment of *in vivo* epithelial barrier function using the FITC-dextran permeability assay showed that MES treatment significantly reduced intestinal permeability at sacrifice, whereas MES+Zn only showed decreasing trend (**Fig. 2f**). Interestingly, along with MES and MES+Zn, treatment with Zn alone reduced fecal MPO (**Fig. 2b, c**), although the underlying mechanism for this effect is beyond the scope of this study.

Next, we tested the effect of MES on the DEX-remission experimental approach using SAMP mice (i.e. established ileitis) (**Fig. 2a**). Remission was confirmed by sacrificing a subset of mice immediately following DEX treatment. The remaining mice then received daily oral gavage of MES or MES+Zn (1 mM) for 50 days. MES+Zn treated group shows significantly higher fecal MES levels at sacrifice confirming that MES+Zn allows MES to reach the colon (**Fig. 2g**). Flow cytometric analysis showed that both MES and MES+Zn reduced RORγt expression in CD4 and CD8 T cells isolated from inguinal lymph nodes (ILN) compared with vehicle-treated controls (**Fig. 2h, i**). Notably, MES+Zn suppressed RORγt more strongly than MES alone. Both MES and MES+Zn reduced fecal MPO levels **(Fig. 2j, k)**, intestinal permeability (*in vivo)* **(Fig. 2l)** and the percentage of abnormal mucosa (3D-SM) at week 7 **(Fig. 2m)** compared to no treatment group. No difference between the groups was observed in ileal histology scores (**Figure S4c, d**). Taken together, these data show that while both MES and MES+Zn relieve intestinal inflammation, MES+Zn provides a greater suppression of the pro-inflammatory RORγt program in SAMP model and maintained higher MES level than MES alone in both models, supporting the use of MES+Zn as a more effective formulation for local immunomodulation.

### Development and characterization of genetically engineered EcN-CAD for enhanced MES production

In the SAMP DEX-remission model, the observed changes in the transcription factors RORγt and FOXP3 in the ILNs, suggest that MES+Zn exerts its therapeutic effects, at least in part, through systemic immunomodulation. To enable sustained intestinal supplementation of MES for local delivery, we sought to engineer a probiotic strain capable of generating MES in situ. As no bacterial enzyme is currently known to directly synthesize MES, *Escherichia coli* Nissle 1917 DE3 (EcN) was engineered to express cis-aconitate decarboxylase (CAD), an enzyme that converts cis-aconitate to ITA, with the hypothesis that ITA could be further converted to MES through endogenous bacterial metabolism. CAD was introduced into EcN via electroporation of the pET-HIS-CAD plasmid, generating the strain of EcN-CAD. Successful plasmid introduction was confirmed by antibiotic selection (**Fig. S3b**).

Following Isopropyl (beta)-D-1-thiogalactopyranoside (IPTG) induction, ITA was not detected in the supernatants of either EcN or EcN-CAD cultures by LC-MS/MS (**Fig. S3a**). In contrast, MES was detected in EcN-CAD cultures, indicating downstream metabolic conversion of CAD-derived intermediates. To directly assess whether EcN-CAD could generate MES from ITA, EcN and EcN-CAD cultures were supplemented with 1 mM ITA and incubated for 24 h without IPTG induction. LC-MS/MS analysis showed significant increase of MES production in the culture supernatant (**Fig. S3c, d**). These results demonstrate that EcN-CAD can produce MES from ITA through bacterial metabolic processes.

Notably, EcN-CAD generated over 10-fold higher MES production than wild-type EcN (**Fig. 3a**; P value < 0.001) in 4 hours after inducing IPTG in M9 broth. To investigate if MES could be continuously released in intestine, IPTG-induced EcN-CAD were added to simulated intestinal fluid (SIF) supplemented with glucose to evaluate MES releasing profile. LC-MS/MS analysis suggests that EcN-CAD released MES in SIF sustained in sustained manner over 24 hours (**Fig. 3b**), indicating the potential for effective intestinal delivery of MES.

**Figure 3:**
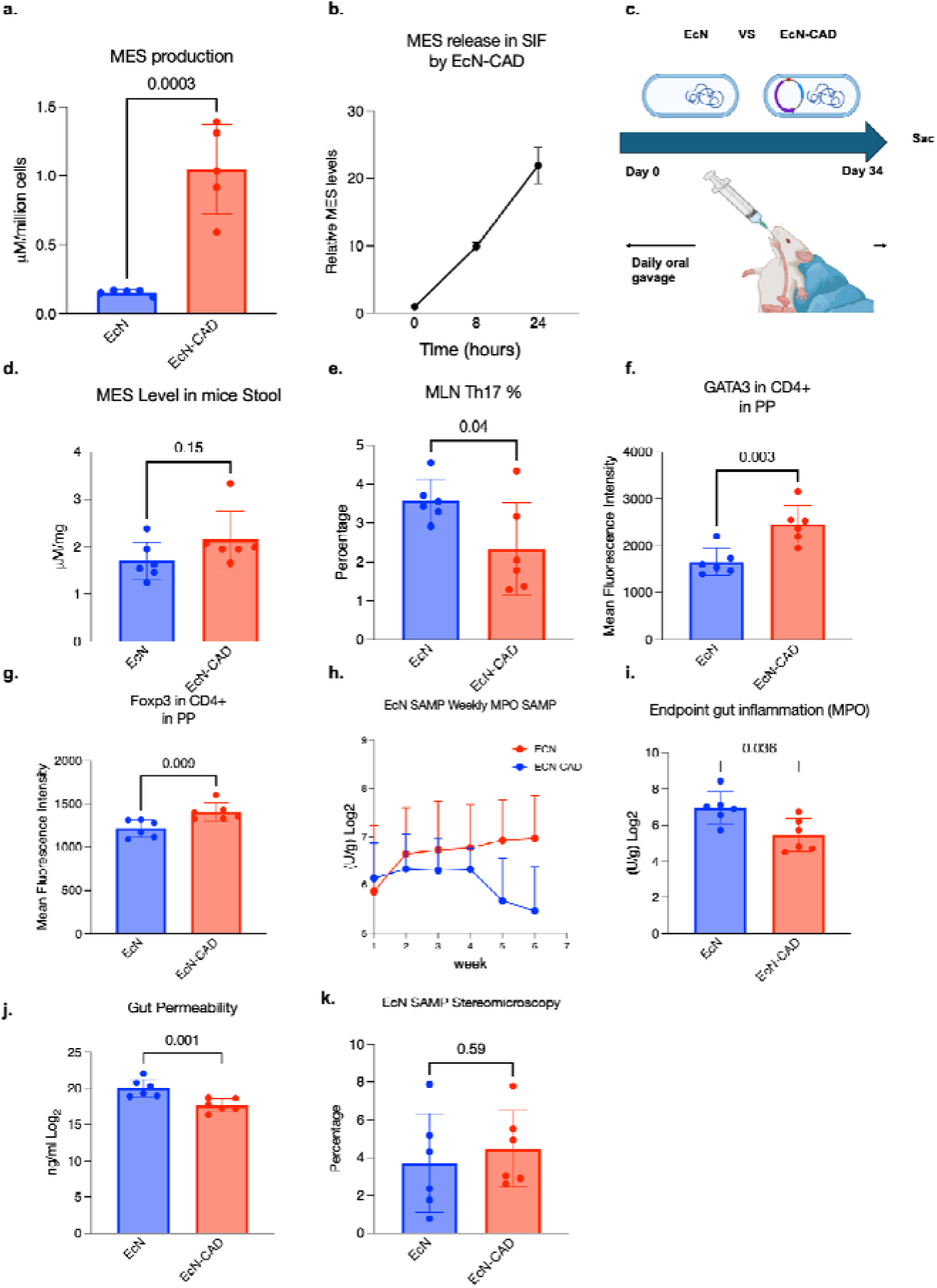
EcN-CAD release MES *in vitro* and decreases inflammatory T cell responses. **(a)** MES production increased by 10-fold in EcN-CAD compared to EcN. Student’s Unpaired two-tailed t-test, n = 4 biological replicates. **(b)** MES release in SIF at 0 hours, 8 hours and 24 hours. N = 6 biological replicates **(c)** 14-week-old SAMP mice were orally gavaged with EcN or EcN-CAD daily for 6 weeks (42 days). **(d)** MES level in mice stool samples at sacrifice. **(e-g)** Flow cytometry analysis of T cell isolated from mesenteric lymph node (MLN) and payers’ patches (PP). **(h-i)** Weekly fecal MPO in SAMP mice over 6 weeks and at week 6 (sacrifice). **(j)** FITC-dextran gut permeability assay measured *in vivo*. **(k)** 3-D Stereomicroscopy of % abnormal mucosa at sacrifice in SAMP mice. P value shows Student’s Unpaired two-tailed t-test, n = 6 mice.

### EcN-CAD reduces pro-inflammatory T cell markers and intestinal inflammation in the SAMP model of CD-ileitis

*In vivo* efficacy of EcN-CAD was assessed in 14-week-old SPF SAMP mice (i.e. established ileitis) by daily oral gavage of EcN-CAD (**Fig. 3c**). EcN-CAD significantly reduced CD8□T cells and T_h_17 population in mesenteric lymph nodes (MLN) (**Fig. 3e**, **Fig. S2l**), while increasing Foxp3 and GATA3 in Peyer’s patches (PP) (**Fig. 3f, g**). EcN-CAD-treated mice also exhibited significantly reduced fecal MPO levels measured weekly and at sacrifice (**Fig. 3h, i**) as well as improved gut barrier function (FITC-dextran) (**Fig. 3j**). However, no difference in abnormal mucosa percentage **(Fig. 3k)** or ileal histology scores (**Fig. S4e, f)** were observed. These data demonstrate that MES-producing EcN-CAD can improve gut permeability, reduce CD gut inflammation, and decrease pro-inflammatory immunophenotype in lymphoid organs when compares to EcN.

### EcN-CAD improves gut barrier integrity in the SAMP mouse remission model

To evaluate therapeutic effects in clinical treatment in CD patients, we again used our DEX remission model to test the ability for EcN-CAD to prolong remission. Following DEX treatment, 14-week-old SAMP mice received daily oral gavage of EcN-CAD or saline vehicle for 4 weeks (**Fig. 4a**). Fecal MES levels were elevated in DEX+EcN-CAD group compared to mice treated + saline group but the difference is not significant (**Fig. 4b**). Compared to DEX+saline controls, DEX+EcN-CAD significantly reduced the frequency of CD8□T cells and RORγt expression in CD4 T cells in mLN (**Fig. 4c, d**). EcN-CAD-treated mice had reduced RORγt expression in both CD4□ and CD8□T cells, and decreased Tbet expression in CD8 T cells within PP (**Fig. 4f-g**). At sacrifice, DEX+EcN-CAD treated mice exhibited significantly reduced gut permeability, but not fecal MPO levels (**Fig. 4h-j**) or histology score (**Fig. S4g, h)** compared to DEX-only control mice. These results suggest that EcN-CAD improve both gut permeability and immunological control in CD.

**Figure 4:**
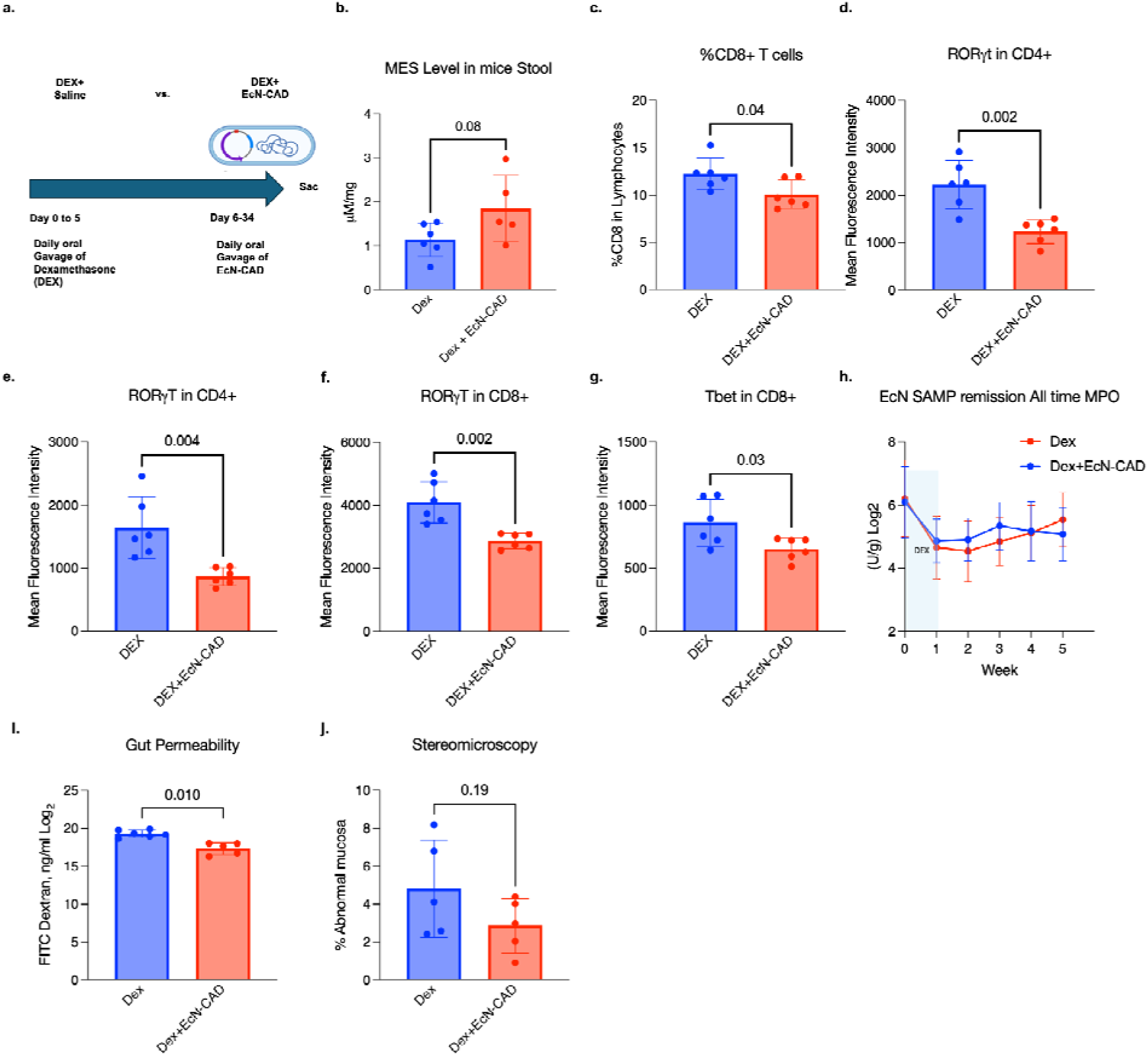
EcN-CAD decrease inflammation in SAMP remission model. **(a)** Schematic of the SAMP-DEX remission model (5-day DEX) followed by EcN-CAD administration for 28 days. **(b)** MES level in mice stool samples at sacrifice. **(c)** Frequency of CD8^+^ T cells in mLN. **(d, e)** gMFI of RORγt expression in CD4^+^ T cells in MLN and PP. **(f)** gMFI of RORγt expression in PP CD8^+^ T cells. **(g)** gMFI of Tbet expression in PP. **(h)** Weekly fecal MPO. **(i)** FITC-dextran *in vivo* gut permeability assay. (**j)** 3-D Stereomicroscopy scores comparing % abnormal mucosa to normal mucosa. P-value shows Student’s Unpaired two-tailed t-test, n = 6 replicates.

### EcN-CAD attenuates ileitis onset in SAMP mice, with lactose enhancing its therapeutic effect

Lactose, a common dietary carbohydrate, activates the lactose operon encoded by the pET-HIS-CAD plasmid, driving high-level CAD expression in EcN-CAD and thereby enhancing intestinal MES production. To determine whether lactose can serve as an effective inducer of CAD expression for sustained MES production, lactose was supplemented to EcN-CAD culture. LC-MS/MS analysis of the supernatant confirmed that lactose significantly increased MES levels in EcN-CAD cultures (**Figure 5a**), suggesting that lactose can enhance EcN-CAD mediated MES release.

**Figure 5:**
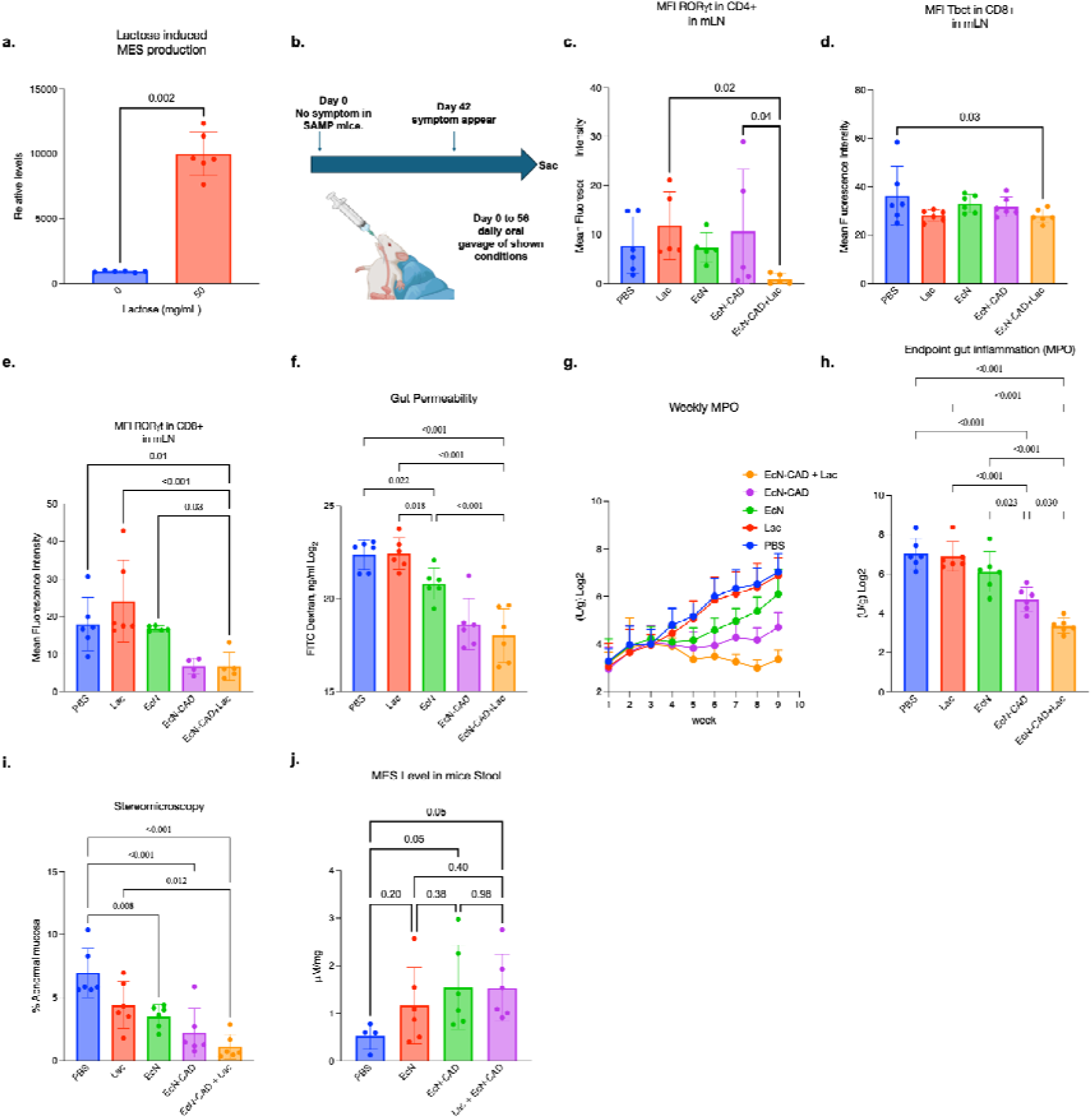
EcN-CAD treatment attenuates ileitis onset in young SAMP model. **(a)** Lactose induces overproduction of MES from EcN-CAD. A 50 mg/mL of lactose were supplemented to LB broth to simulate lactose concentration in milk. **(b)** Four –week-old SAMP mice were orally gavaged with saline or Lac or EcN or EcN-CAD or EcN-CAD+Lac for 8 weeks (56 days). **(c)** gMFI of RORgt expression in CD4^+^ T cells in MLN. **(d)** gMFI of Tbet expression in the CD8^+^ T in MLN. **(e)** gMFI of RORgt expression in CD8^+^ T cells in MLN. **(f)** FITC *in vivo* gut permeability. **(g, h)** Weekly fecal MPO in SAMP mice over 8 weeks and at week 8 (sacrifice). **(i)** Scores from 3-D Stereomicroscopy. **(j)** MES level in mice stool samples at sacrifice. n = 5-6 biological replicates, One-way ANOVA, Fisher’s LSD post-hoc test; P-value between EcN-CAD+Lac and other groups.

To determine whether EcN-CAD attenuates ileitis onset, 4-week-old SAMP mice (i.e., before CD-ileitis onset) were orally gavaged daily for 8 weeks with either EcN-CAD alone or EcN-CAD supplemented with lactose (EcN-CAD+Lac). Control groups received PBS, lactose alone, or wild-type EcN (**Fig. 5b**). On day 56, mice were sacrificed, and MLN cells were harvested and analysed by flow cytometry for expression of CD4, CD8, Tbet, RORγt, GATA3, Foxp3, CD44, and CD25. EcN-CAD+Lac treatment significantly reduced RORγt expression in CD4□ T cells compared to EcN, lactose, and PBS controls (**Fig. 5c**). Additionally, Tbet expression in CD8 T cells and RORγt expression in CD8 T cells were significantly lower in the EcN-CAD+Lac group compared to the lactose and EcN-CAD groups (**Fig. 5d, e**). Importantly, EcN-CAD significantly improved gut barrier function and reduced fecal MPO compared with PBS-, lactose– and EcN-treated mice, and these improvements were further enhanced in the EcN-CAD+Lac group (**Fig. 5f– h**). Lactose supplementation provided additional benefit on T-cell phenotype, significantly reducing CD8 Tbet and CD4 RORγt expression relative to EcN-CAD alone (**Fig. 5d, e**). Lactose supplementation had an incremental effect on intestinal permeability over EcN-CAD alone (**Fig. 5f**), consistent with near-maximal barrier restoration by EcN-CAD. No difference was seen in spleen T_reg_ percentage, Th_17_ and gMFI of RORγt in EcN-CAD+Lac group compared to all other treatment groups, suggesting no systemic change was observed in splenic T cell subsets. Also, EcN-CAD group significantly increase T_reg_ percentage in spleen when comparing to EcN group (**Fig. S3a**). Additionally, 3D-stereomicroscopic assessment of ileal tissues showed significantly lower %abnormal mucosa in both EcN-CAD and EcN-CAD+Lac groups when compared with control mice (**Fig. 5i**). In EcN-CAD and EcN-CAD+Lac group, significant upregulation of MES in mice stool samples were observed when compared to PBS treated group (**Fig. 5j**). Interestingly, only EcN-CAD treated group shows significant decrease in histology score compared to PBS treated control group (**Fig. S4i, j**). Together, these findings suggest that lactose-inducible EcN-CAD could improve gut barrier function as well as modulate pro-inflammatory T cell phenotypes in a preclinical model of CD-ileitis.

## Discussion

This study identifies MES as a promising therapeutic metabolite for controlling intestinal inflammation in Crohn’s disease (CD). Fecal MES levels correlated with mucosal inflammatory burden in CD patients, linking MES to disease-associated inflammation. *In vitro*, MES suppressed Th17 differentiation, with proteomic analysis implicating OXPHOS and mitochondrial pathways as potential mediators. *In vivo*, restoring local MES levels through either an engineered probiotic or a MES+Zn formulation reduced Th17-associated intestinal inflammation across therapeutic, remission-maintenance, and prophylactic settings in mice prone to CD-like ileitis. Together, these findings support local MES restoration as a potential therapeutic strategy for CD.

In CD patients, fecal MES levels correlated with fecal MPO (Pearson r = 0.47), a validated surrogate of neutrophil infiltration and epithelial injury^23^, suggesting that MES is actively produced at sites of mucosal inflammation and may reflect a local immunoregulatory response to injury. This aligns with evidence that the fecal and serum metabolome in CD is remodeled during disease activity and may predict future flares^24^. These data also align with the broader concept that barrier disruption and neutrophil activation drive disease-associated metabolic shifts. Critically, this association was specific to MES and not its isomer ITA, indicating a non-redundant, IBD-associated role for MES as a candidate biomarker and therapeutic target in CD.

The correlation between fecal MES and mucosal inflammation raised the question of whether restoring MES could counteract the RORγt program driving CD pathology. MES significantly suppressed RORγt expression in activated mouse CD3^+^ T cells *in vitro*, demonstrating for the first time that MES can suppress the Th17 axis in adaptive immune cells. Aso et al. showed that ITA itself ameliorates autoimmunity through metabolic and epigenetic T-cell reprogramming in mice, and our results suggest MES may do the same. Th17 cells are pathologically expanded in the CD intestinal mucosa and required for disease progression, however pharmacological targeting of the Th17 axis via anti-IL-17A agents has shown limited clinical benefit and can worsen the disease condition^4,5,10^. Rather than neutralizing a single downstream cytokine, MES suppresses the upstream RORγt-associated inflammatory program and T-cell metabolic activity, positioning MES as a candidate immunometabolic regulator of pathogenic T-cell responses in CD.

Global proteomics of MES-treated T cells identified broad suppression of OXPHOS and mitochondrial respiratory complex proteins, suggesting MES acts partly through metabolic rewiring. He et al. found that MES modulates macrophage inflammatory responses without affecting mitochondrial function or TCA cycle activity in that myeloid lineage, distinguishing MES from ITA. The OXPHOS suppression we observe in T cells therefore likely reflects a cell-type-specific effect, consistent with the known divergence in mitochondrial metabolism between innate and adaptive immune cells^25,26^. Whether this metabolic suppression is causal to Th17 inhibition, or a parallel response, will require targeted genetic validation.

The *in vitro* evidence indicates that MES suppresses RORγt expression and OXPHOS and mitochondrial activity supported directly testing whether exogenous MES delivery could reduce intestinal inflammation in two genetically distinct mouse models of CD-ileitis. MES+Zn reduced inflammation and improved 3D stereomicroscopy abnormal mucosal percentage in both the TNFΔARE and SAMP remission models, suggesting pharmacological MES delivery. Because mucosal healing proceeds through a defined sequence, improvements in permeability, fecal MPO, and macroscopic disease symptoms precede histological recovery^27–31^. Permeability and fecal MPO track early restoration of tight junction integrity and resolution of acute inflammation, whereas histology reflects the slower reversal of microscopic damage such as immune cell infiltration and crypt injury^32–34^. The two readouts also differ in sampling, i.e. 3D-StereoMicroscopy assessment surveys the entire colon and ileum, whereas histology scores survey discrete tissue sections, and this difference in coverage further accounts for the inconsistency between macroscopic and histological findings^35^. We expect that with longer treatment protocols these indications of the disease will improve as well.

The enhanced mucosal bioavailability of MES+Zn is consistent with zinc-chelation strategies used in colonic-targeted nanomedicines, where charge-mediated retention at the inflamed epithelium enhances local drug concentration^17,36^. MES+Zn prolonged remission in both models, which suggests metabolite restoration alone may be sufficient to attenuate intestinal inflammation, supporting sustained luminal MES as a therapeutic strategy. However, the RORγt suppression observed in the ILN indicates that MES and MES+Zn also act systemically, which motivated us to develop a platform for localized delivery.

The *in vivo* efficacy of MES supplementation motivated development of a microbial delivery strategy, given that the gut microbiome is a major determinant of mucosal immune homeostasis and a tractable therapeutic target in CD^37,38^. Naturally occurring probiotics such as *Lactococcus lactis* and *Escherichia coli Nissle* 1917 (EcN) have long been explored as live biotherapeutics for CD, and engineered variants delivering anti-inflammatory cytokines or enzymes have shown preclinical efficacy^32–35^. Though clinical translation has been limited by the difficulty of sustaining therapeutic concentrations at the disease site. Moreover, conventional probiotic supplementation with defined strains such as *Lactobacillus rhamnosus* GG has shown limited, inconsistent efficacy in CD^39,40^. EcN-CAD addresses these limitations by continuously delivering a single, chemically defined immunometabolite directly at the intestinal mucosa. It extends earlier engineered-probiotic approaches, including IL-10-secreting *L. lactis*^41^, anti-TNFα nanobody platforms^42^ and programmable cytokine circuits^43^ by using the bacterium’s own biosynthetic machinery to produce MES. Importantly, EcN-CAD also introduces dietary tunability, allowing biotherapeutic output to be adjusted in real time through a non-pharmacological approach.

The therapeutic and maintenance efficacy of EcN-CAD raised the question of whether pre-emptive luminal MES delivery could prevent disease onset altogether. EcN-CAD prevented intestinal inflammation in pre-ileitis SAMP mice, an effect further enhanced by dietary lactose supplementation. This dietary tunability distinguishes EcN-CAD from conventional engineered probiotics and existing biologics and is conceptually related to microbiome-modulating dietary strategies demonstrated in other inflammatory contexts such as food allergy. Preventive efficacy is especially relevant in CD because high-risk individuals can increasingly be identified before clinical inflammation develops. Genetic risk assessment, biomarkers, and endoscopic surveillance may help define this early intervention window^8,9,24^ EcN-CAD could be particularly useful in this setting. Its activity is self-limiting and can potentially be adjusted through diet.

Although this study demonstrates a correlation between MPO levels and MES levels, as well as the therapeutic benefits of local MES delivery in reducing intestinal inflammation and disease symptoms, several questions remain unanswered. Whether fecal MES depletion drives or results from mucosal inflammation is unclear; longitudinal monitoring in prospective CD cohorts could establish whether declining MES precedes flares and serves as a non-invasive biomarker of remission^24^. The proteomic candidate mechanisms, actin remodeling, mitochondria and OXPHOS suppression, require genetic and pharmacological validation, and extending the analysis to lamina propria T cells, macrophages, and epithelial cells will clarify the full scope of MES activity. The efficacy of EcN-CAD in SAMP and MES+Zn in TNFΔARE models warrants evaluation in additional preclinical systems and, ultimately, human trials^2,9^. Clinical development will further require defining achievable luminal MES pharmacokinetics in humans, biocontainment assessment of the engineered strain, and evaluation of its colonization dynamics and impact on the endogenous microbiome. The bacterial strain’s antibiotic-resistance marker also limits translation; an antibiotic-free version and an inducer alternative to lactose for lactose-intolerant patients will be important next steps.

In summary, this work identifies MES as a CD-relevant immunometabolite whose restoration at the intestinal mucosa suppresses Th17-driven inflammation, improves gut barrier integrity, and prevents disease onset in preclinical models. Proteomic analysis points to mitochondrial function suppression as a candidate mechanism underlying MES-mediated Th17 inhibition, advancing our understanding of how mesaconic acid may differentially shape immune responses across cell lineages^20,21,26,44^. The EcN-CAD platform translates this biology into a self-sustaining, dietarily tunable delivery system that achieves localized mucosal MES concentrations. This approach addressing both the limitations of systemic immunosuppression and the insufficient endogenous MES levels in CD patients. More broadly, this work supports restoring endogenous immunometabolic pathways via engineered living therapeutics as a tractable strategy for mucosal immune homeostasis in CD, with potential relevance to other inflammatory diseases.

## Supplementary Figures

**Figure S1.**
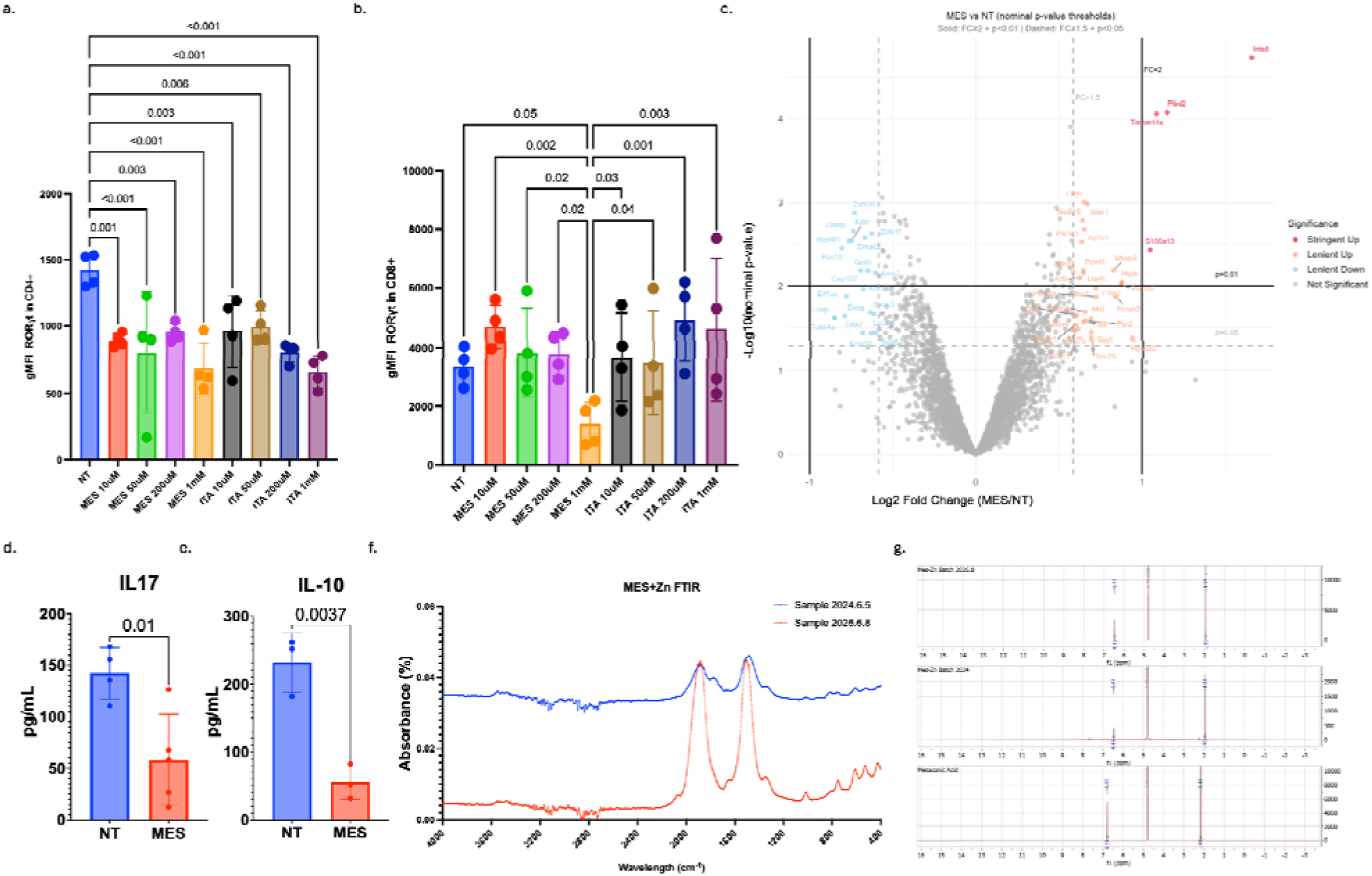
MES and MES+Zn effects on mice and human T-cell markers, and cytokine concentration in supernatant. (a-b) RORgt geometric mean fluorescence intensity (gMFI) of CD4 and CD8 T cells when mouse CD3^+^ T cells were treated with different concentration of MES and ITA. One-way ANOVA; n = 4 mice. (c) Volcano plot of mouse CD3^+^ T cell global proteomics after treating with MES vs NT. One-way ANOVA; n = 4 mice. (d-e) IL-17 and IL-10 ELISA in the supernatant of activated mouse CD3^+^ T cells treated with MES vs NT. One-way ANOVA; n = 4 mice. (f-g) FTIR and NMR of MES+Zn for sample synthesized in 06/05/2024 and 06/08/2026.

**Figure S2.**
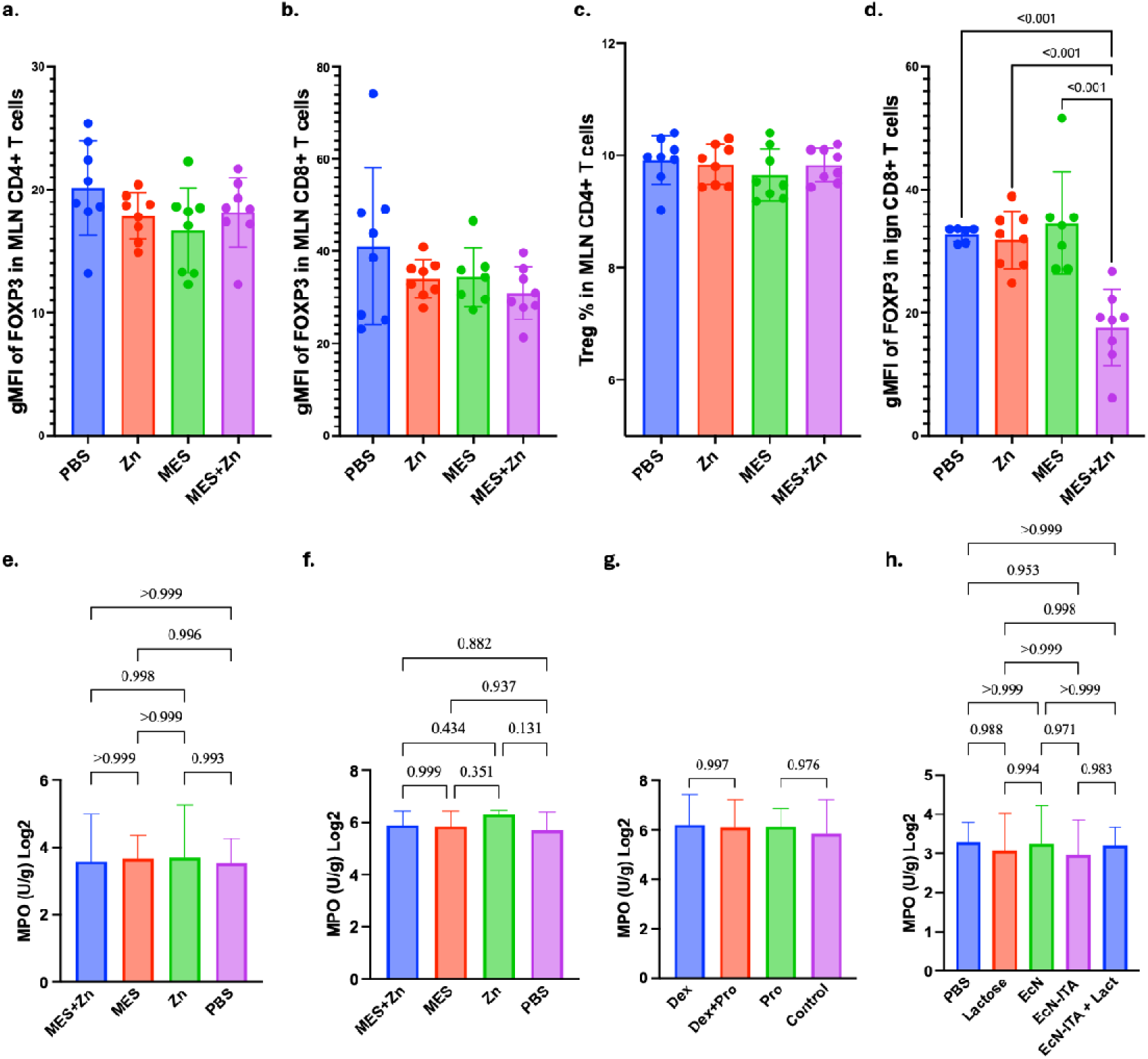
MES production from itaconate by EcN-CAD, and additional in vitro and mucosal analyses. (a-d) Flow cytometry analysis of FOXP3 gMFI in CD4^+^ and CD8^+^ T cells in MLN and ILN. One-way ANOVA; n = 8 mice. (e) Base fecal MPO of SAMP mice for MES+Zn SAMP remission model. One-way ANOVA; n = 8 mice. (f) Base fecal MPO of TNFDARE mice that treated with MES+Zn. One-way ANOVA; n = 6 mice. (g) Base fecal MPO of SAMP mice for SAMP and SAMP remission model that treated with EcN-ACD. One-way ANOVA; n = 6 mice. (h) Base fecal MPO of SAMP mice for young SAMP model that treated with EcN-ACD. One-way ANOVA; n = 6 mice.

**Figure S3.**
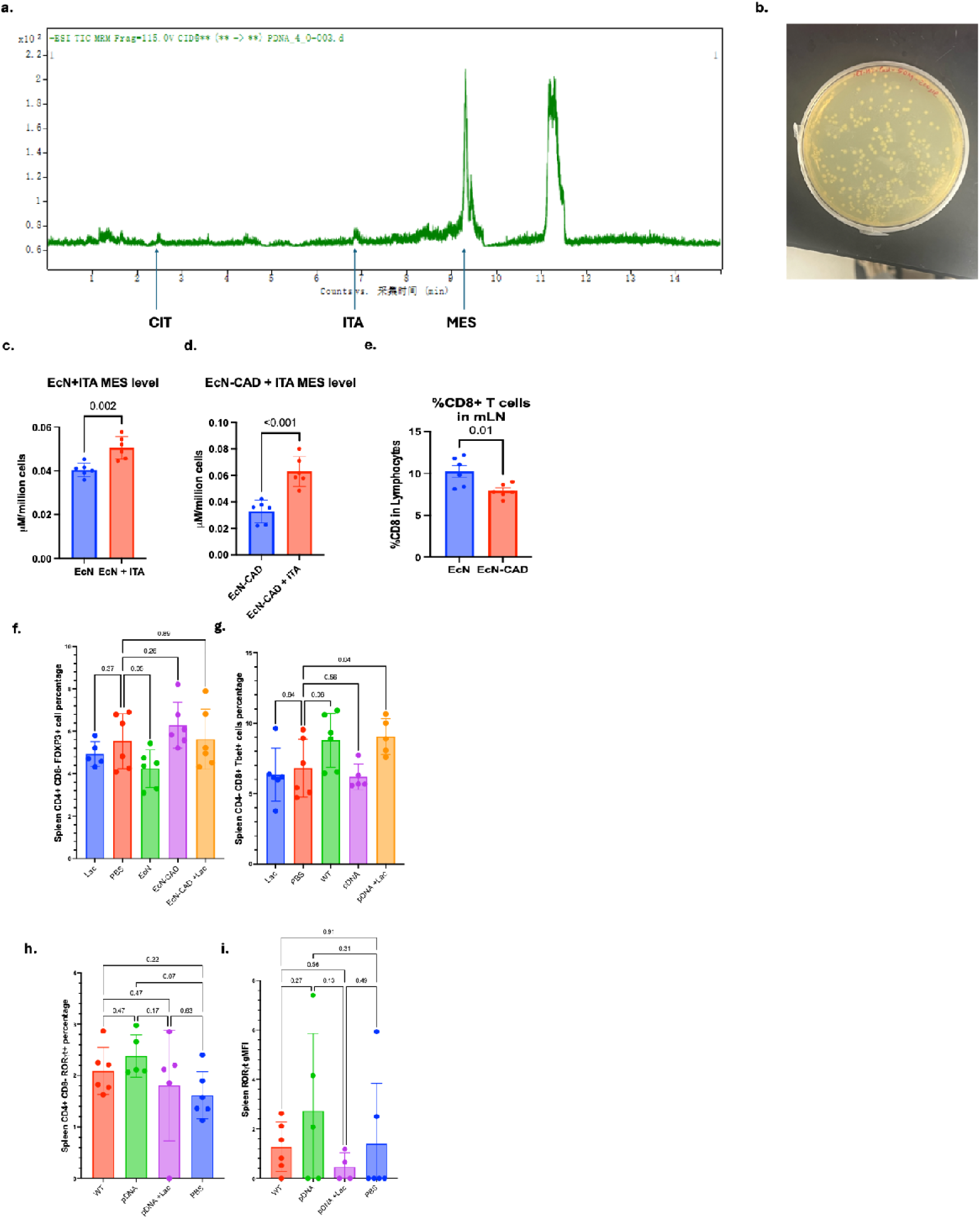
(a) Representative data of LC/MS-MS of EcN-CAD treated with IPTG supernatant. (b) Image of EcN-CAD colonies on ampicillin ager plate. (c–d) MES level evaluated by LC-MS/MS in EcN&EcN-CAD supernatant w/wo 1mM ITA supplement. Two-way unpaired two tailed student’s T test, n = 6 replicates. (e) Flow cytometry analysed CD8 % in MLN. Two-way unpaired two tailed student’s T test, n = 6 mice. (f-i) Flow cytometry analysed of T cells in spleen in young SAMP model. One-way Anova; n = 6 mice.

**Figure S4.**
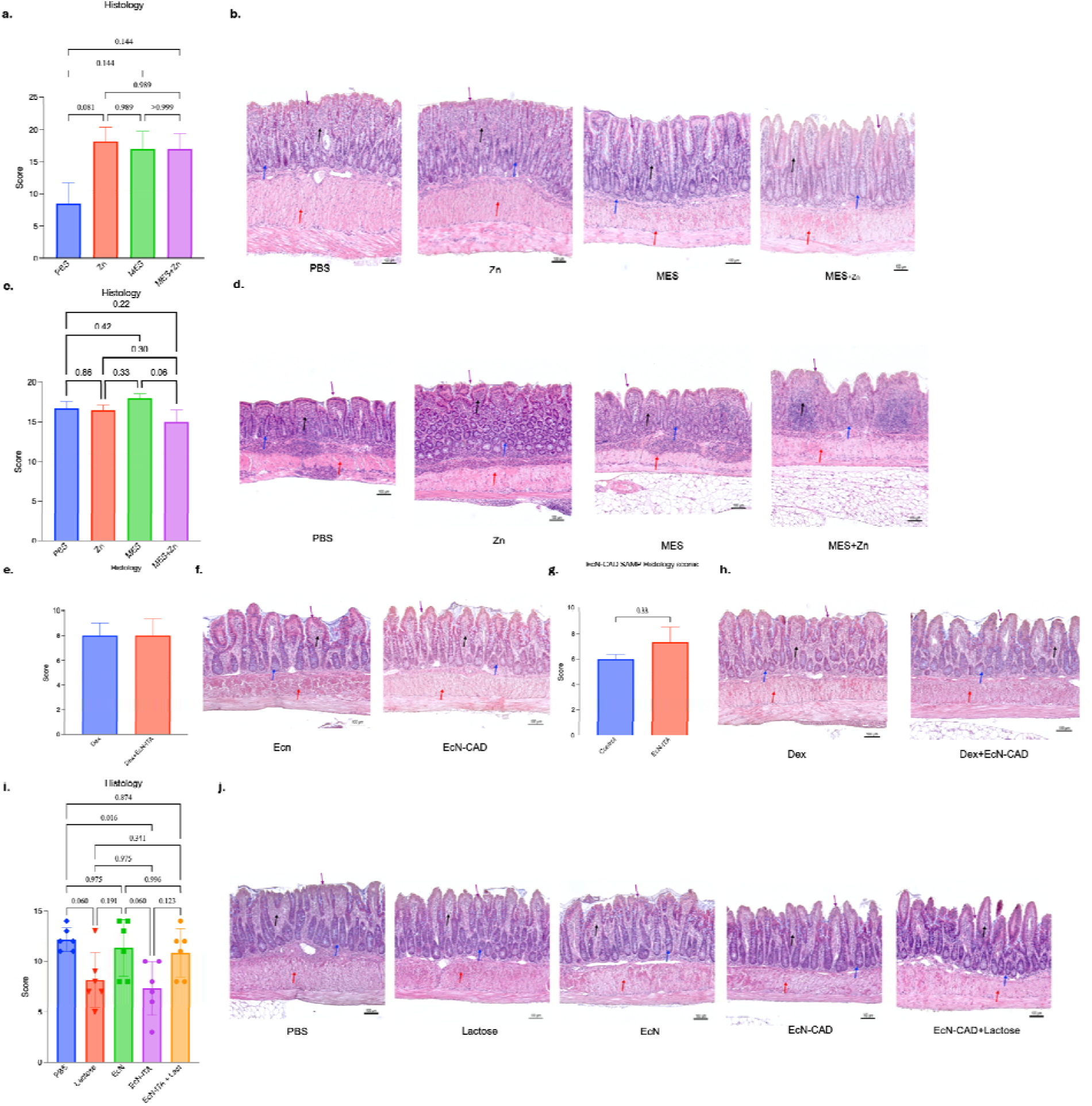
Histological score and representative H&E staining sections of all *in vivo* studies. (a) Histology score of TNFΔARE mice. One-way Anova, n = 6 mice. (b) Representative histopathological sections of H&E-stained ileal tissues from TNFΔARE mice showing an increase in the cellular infiltration (black arrows), mucosal thickening (red arrows), fused and distorted villi architecture depicting altered epithelial structure (purple arrows) and elongation of the crypts (blue arrows) in the PBS and Zn treated mice, however, the treatment with MES and MES+Zn show decreased cellular infiltration (black arrows), preservation of villi architecture (purple arrows) lesser crypt elongation (blue arrows) and diminished thickening of the muscularis mucosa (red arrows). (c) Histology score of SAMP mice. One-way Anova, n = 8 mice. (d) Representative histopathological sections of H&E-stained ileal tissues from SAMP mice showing an increase in the cellular infiltration (black arrows), mucosal thickening (red arrows), fused and distorted villi architecture depicting altered epithelial structure (purple arrows) and elongation of the crypts (blue arrows) in the PBS and Zn treated mice, however, the treatment with MES and MES+Zn show decreased cellular infiltration (black arrows), preservation of villi architecture (purple arrows) lesser crypt elongation (blue arrows) and diminished thickening of the muscularis mucosa (red arrows). (e) Histology score of SAMP mice. One-way Anova, n = 6 mice. (f) Representative histopathological sections of H&E-stained ileal tissues from SPF-SAMP mice showing similar cellular infiltration (black arrows), altered epithelial structure (purple arrows), crypt structure (blue arrows) and mucosal thickening (red arrows) in the EcN and EcN-CAD treated mice. (g) Histology score of SAMP mice. One-way Anova, n = 6 mice. (h) Representative histopathological sections of H&E-stained ileal tissues from SPF-SAMP mice showing similar cellular infiltration (black arrows), altered epithelial structure (purple arrows), crypt structure (blue arrows) and mucosal thickening (red arrows) in the EcN and EcN-CAD treated mice. (a) Histology score of SAMP mice. One-way Anova, n = 6 mice. (j) Representative histopathological sections of H&E-stained SAMP ileal tissues show increased in the cellular infiltration (black arrows) and mucosal thickening (red arrows). Fused and distorted villi architecture depicting altered epithelial structure (purple arrows<u>). E</u>longation of the crypts (blue arrows) shown (blue arrows) in the PBS-, Lactose<u>-</u>and EcN-treated mice. Note that however, the treatment with EcN-CAD and EcN-CAD+Lactose treated group show decreased cellular infiltration (black arrows), preservation of villi architecture (purple arrows) lesser crypt elongation (blue arrows) and diminished thickening of the muscularis mucosa (red arrows). *Hyperproliferation of goblet cells in PBS treated mice.

**Figure S5.**
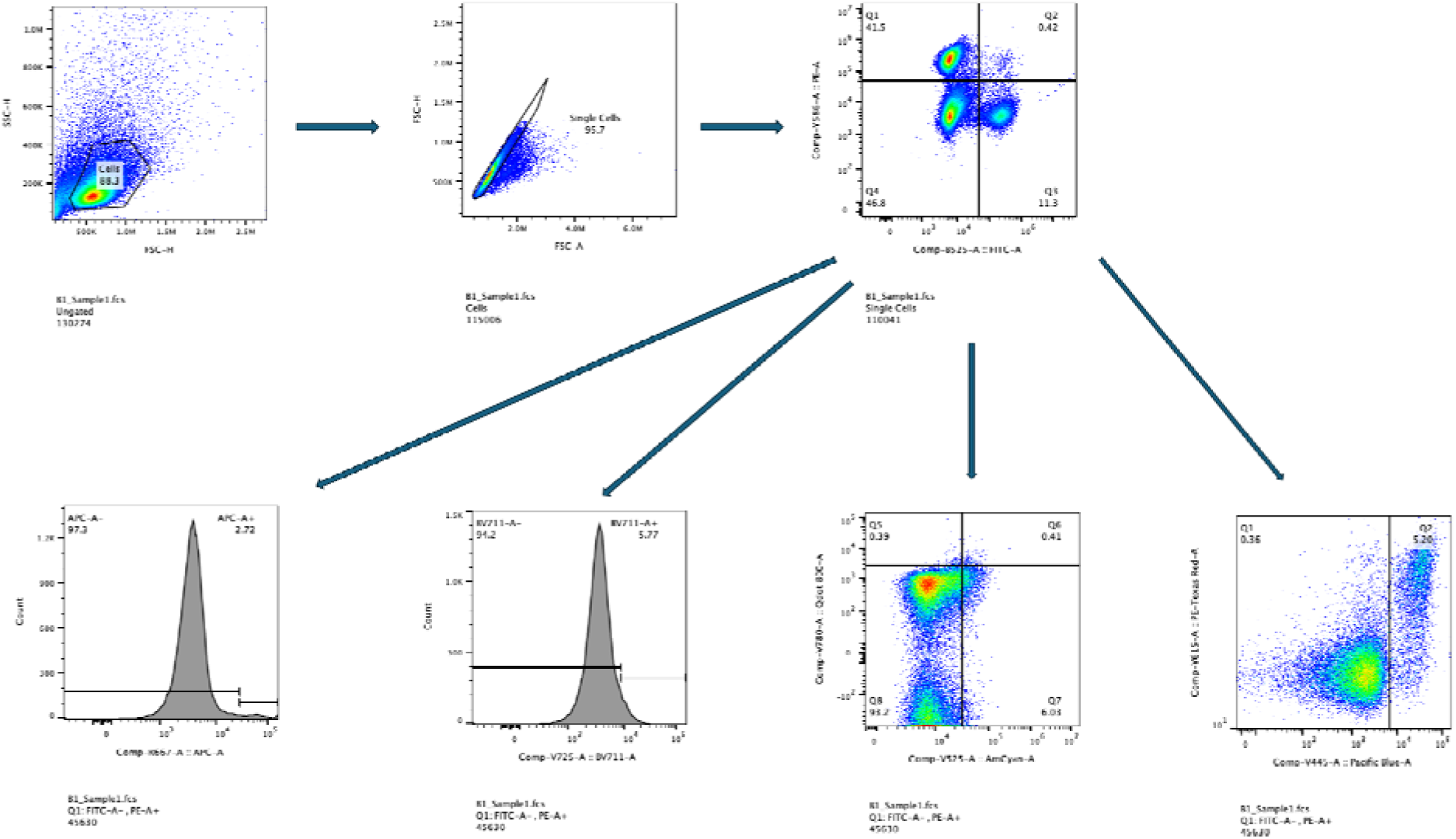
Gating strategy for flow cytometry.

**Figure S6.**
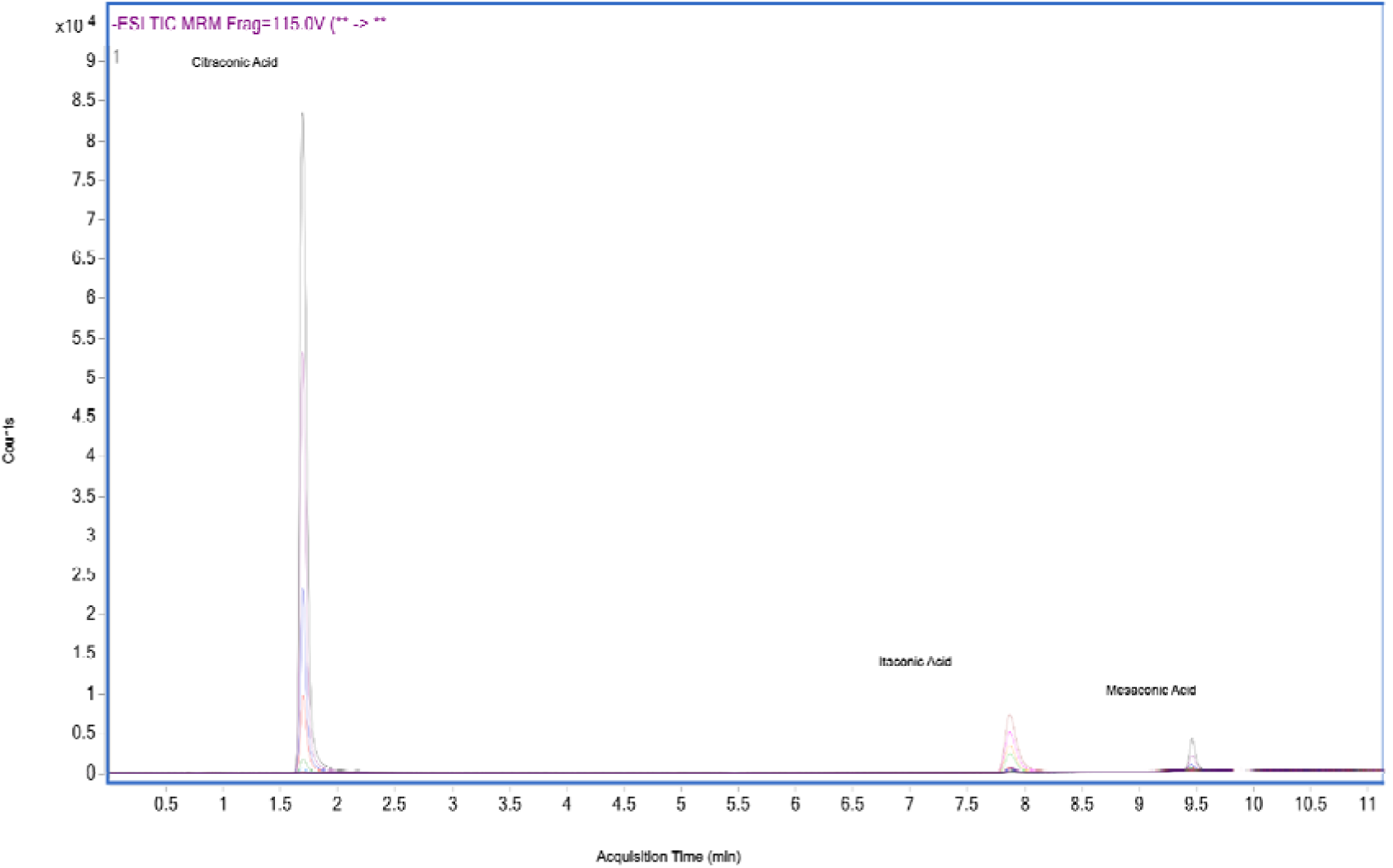
Representative ion chromatograms of standards for itaconic acid, citraconic acid, and mesaconic acid. TIC MRM counts for itaconic acid (1, 10, 20, 50, 100 µmol/L), citraconic acid (1, 10, 20, 50, 100 µmol/L), and mesaconic acid (1, 10, 20, 50, 100 µmol/L) vs. acquisition time.

## Methods and Materials

### MES+Zn Synthesis

Mesaconate–zinc (MES+Zn) formulations were prepared by dissolving mesaconic acid, zinc chloride (ZnCl□), and sodium hydroxide (NaOH) in deionized (DI) water. Briefly, mesaconic acid (1 g), ZnCl□(1 g), and NaOH (1.02 g) were combined in a 15 mL conical tube, followed by the addition of 13 mL DI water. The mixture was vortexed until complete dissolution was achieved. The pH of the solution was adjusted to 7.4 by dropwise addition of 22% (w/v) NaOH under gentle mixing. An aliquot (250 µL) of the resulting solution was transferred into a 50 mL conical tube and diluted with 30 mL of 10× phosphate-buffered saline (PBS) to generate the working MES+Zn formulation.

### Bacterial Strains, Culture, and Induction

*Escherichia coli* Nissle 1917(DE3) Drnc (EcN) was obtained from Creative Biolabs. All bacterial cultures were maintained in LB broth at 37 °C with shaking at 200 rpm. Single colonies were selected from LB agar plates containing 100 mg/mL ampicillin (Sigma-Aldrich).

For IPTG-based induction, bacterial overnight cultures were 1:100 diluted in fresh LB broth and were grown to an optical density at 600 nm (OD□□□) of 0.6, then resuspended in M9 minimal medium supplemented with 0.8 mM isopropyl β-D-1-thiogalactopyranoside (IPTG) (Sigma Aldrich & ThermoFisher) and incubated for 4 h or 24 h. A total of 50 mg /mL of ampicillin was supplemented to broth during all the process. Induced bacteria were washed three times with phosphate-buffered saline (PBS), resuspended in PBS containing 50% glycerol, and stored at – 80 °C. For lactose induction, lactose was added to 1:100 diluted overnight cultures after OD reached 0.6 to a final concentration of 50 mg/mL and incubated for 24 hours.

### Plasmid Construction and Transformation

pET-His-CAD plasmid is a gift from Dr. Yuji Aso’s lab. Plasmid transformation into EcN was performed by electroporation using a 2-mm gap cuvette and a single 2.5 kV pulse. Transformed bacteria (EcN-CAD) were recovered in SOC medium (ThermoFisher) at 37 °C for 1 h, plated on LB agar containing 100 ng/mL ampicillin.

### Metabolites Methanol Extraction

Methanol (1 mL) was added to 200 µL of each sample and mixed thoroughly. Samples were incubated on an orbital shaker with dry ice for 4 hours, followed by centrifugation at 16,000 rpm. The resulting supernatant was collected, and methanol was evaporated using a vacuum centrifuge. The dried pellets were subsequently resuspended in ultrapure water. The relative concentration of MES in the samples was determined using LC-MS/MS analysis.

### CD3 T Cell Isolation

CD3 T cells were isolated from the spleens of C57BL/6J mice spleen or human PBMC (peripheral blood mononuclear cell) using magnetic bead separation (Miltenyi Biotec). Cells were seeded at 1 × 10 cells/mL in RPMI 1640 medium supplemented with 10% fetal bovine serum (FBS), 1% penicillin–streptomycin and 500 ng/mL CD28 antibody, in CD3 antibody pre-coated plate, and incubated at 37 °C with 5% CO for 72 hours.

### Seahorse Extracellular Flux Assay

CD3 T cells isolated from C57BL/6J spleens were activated with plate-bound anti-CD3 and soluble anti-CD28 for 72 hours, with 1 mM mesaconate or vehicle present throughout. Cells were washed three times in Seahorse XF DMEM (pH 7.4) supplemented with 2 mM L-glutamine and 1 mM sodium pyruvate, containing 10 mM glucose for the Mito Stress Test or no glucose for the Glycolysis Stress Test, counted, and seeded at 2 × 10 cells per well on Poly-D-lysin coated XF96 microplates. Plates were equilibrated for 60 min at 37 °C in a non-CO incubator. Following three baseline measurements, oligomycin (1 mM), FCCP (1.1 mM) and rotenone/antimycin (200 nM) A were injected sequentially to measure OCR; glucose (1 mM), oligomycin (1 mM) and 2-deoxy-D-glucose (10 mM) were injected sequentially to measure ECAR, with three measurements recorded after each injection. Immediately following each assay, cells were stained with Hoechst 33342 and all OCR and ECAR values normalised to DNA content per well. Data were acquired and analysed using Wave software (Agilent). Technical replicates were averaged within each animal, giving n = 4 mice.

### SAMP Mouse Models

SAMP1/YitFc (SAMP) mice were obtained from the Cleveland Digestive Diseases Research Center Mouse Models Core of Case Western Reserve University to evaluate the immunomodulatory effects of EcN-CAD. Mice were maintained under specific pathogen-free (SPF) conditions in germ-free NesTiso cages with non-edible aspen bedding to avoid bedding ingestion ^45^. All mice were fed autoclaved chow (Labdiet® Rat/Mouse 18% VacPac-5LQ6) and sterile water *ad libitum and* were kept under 12-hour light/dark cycle in AAALAC-accredited Animal Research Centre rooms regulated for the temperature and humidity to maintain standard conditions. All animal protocols were approved by the Institutional Animal Care and Use Committee (IACUC protocol 2023-0011) and adhered to AAALAC guidelines. An inter-subject pre-experimental fecal homogenization (IsPreFeh) was performed one week before the start of all experiments to restrict cage-to-cage microbiome variability ^35,46^. n = 6 mice for EcN-CAD experiment and n = 8 mice for MES+Zn SAMP1/YitFc dexamethasone-).

In the established disease model, 14-week-old SAMP mice were orally gavaged daily with either IPTG induced EcN or EcN-CAD (10 CFU in 200 μL of saline) for 28 days. Intestinal barrier function was evaluated using FITC-dextran (4 kDa, 600 mg/kg, orally), and plasma fluorescence was measured 4 hours post-gavage. Fecal myeloperoxidase (MPO) levels were assessed by ELISA. At sacrifice, mesenteric lymph nodes (mLN) and Peyer’s patches (PP) were harvested and processed for flow cytometric analysis.

In the remission model (SAMP-DEX), 14-week-old SAMP mice received intraperitoneal injections of dexamethasone (DEX; 1 mg/kg/day) for 5 consecutive days to induce remission. Mice were subsequently treated with daily oral gavage of saline or IPTG induced EcN-CAD (10 CFU in 200 μL of saline) for 28 days. Gut permeability and fecal MPO were assessed, and lymphoid tissues were collected for immunophenotyping.

In the early-intervention model, 4-week-old SAMP mice (prior to disease onset) were treated for 56 days with daily oral gavage of saline, lactose (10 mg/day), EcN, EcN-CAD, or EcN-CAD combined with lactose (EcN-CAD+Lac). At the study endpoint, FITC-dextran and fecal MPO assays were performed, and mLN were analysed by flow cytometry.

### Histology Assay

The ileal tissue samples were washed in sterile PBS after removal, and were opened longitudinally and fixed in 10% buffered formalin for 24 hours at 4° C. The tissues were rinsed with 70% ethanol before paraffin embedding and were then subjected to Hematoxylin and Eosin (H&E) staining. The degree of inflammation was evaluated by using a validated scoring system using three essential parameters: 1) active inflammation, 2) chronic inflammation, and 3) villus architecture, on a scale of 0 (normal tissue) to 3 (severe inflammation) ^47,48^.

### Assessment of Intestinal inflammation and permeability

Fecal samples were collected weekly to measure the myeloperoxidase (MPO) activity by following a previously described dianisidine-H_2_O_2_ based assay ^49–51^. Tissue specimens were weighed, diluted 20-fold (wt/vol) in 0.5% hexadecyltrimethylammonium bromide (HTAB) buffer, homogenized with a Polytron PT 10-35 tissue homogenizer, and centrifuged to pellet debris. Supernatant (7 µL) was loaded in triplicate into a 96-well plate and mixed with 200 µL of substrate solution (5 mM potassium phosphate buffer containing 0.0005% H_2_O_2_ and 0.167 mg/mL o-dianisidine). The average rate of change in absorbance at 450 nm over two sequential 30-s intervals was used to calculate MPO activity. The intestinal permeability was assessed in all experimental groups before and after the treatment by following a previously described method using a fluorescently labelled sugar probe, fluorescein isothiocyanate (FITC) Dextran^52^. Mice were deprived of food for 4 h and then given 200 µL of 4 kDa FITC-dextran (80 mg/mL; Sigma-Aldrich) by oral gavage. Food and water were withheld for 4 hours after gavage. Blood was then collected by retro-orbital bleeding, and fluorescence intensity was measured in a fluorescence plate reader at an excitation wavelength of 493 nm and an emission wavelength of 518 nm. Higher fluorescence was taken to indicate increased translocation of FITC-dextran from the gut to the circulation and therefore greater intestinal permeability.

### LC-MS Analysis on itaconic acid, citraconic acid and mesaconic acid

Agilent® prescribed settings (Table 1 from published Agilent application notes^53^) were used to determine instrument operating parameters. 1 μL of each sample and standards were injected into a 10 cm long, 2.1 mm i.d., 2.7 μm pore size Agilent Infinity Lab Poroshell 120 HILIC-Z PEEK lined column (Agilent Technologies, CA, United States) for methylmalonic acid separation at a constant flow rate of 0.5 mL/min over 15 min using a multisegmented linear gradient of mobile phase A (10 mM ammonium acetate solution in water with 2.5 µM medronic acid at pH = 9) and B (10 mM aqueous ammonium acetate in water/acetonitrile 10:90 (v:v) with 2.5 µM medronic acid at pH = 9) as follows: 0-11.5 min, 0%-30% A and 100%-70% B; 11.5-12 min, 30%-0% A and 70%-100% B; isocratic period, 12-15 min, 0% A, 100% B. The 15 min sample run is followed by a post-run rinse of 5 min. Each sample was analyzed in triplicate. The mass spectrometer was set as Multiple Reaction Monitoring (MRM) scan type using an AJS source with the following conditions: gas temperature 200; gas flow 10.0 L/min; nebulizer 40 psi; sheath gas temperature 300; sheath gas flow 12 L/min; capillary voltage 3,000 V; nozzle voltage 0 V **(Table 1)**. Mass spectrometer parameters **(Table 2)** such as precursor ion, product ion, collision energy and fragmentor were set according to the work from Katharina et al^54^.

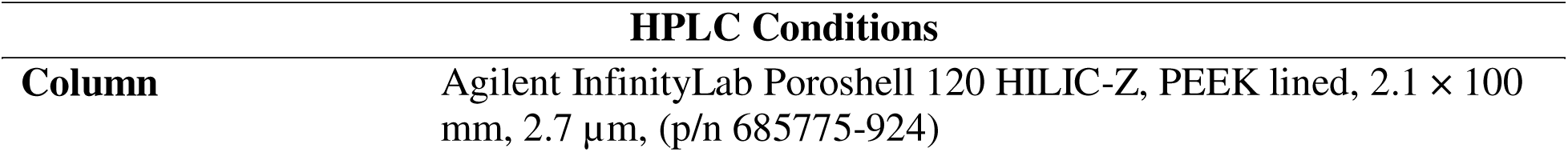
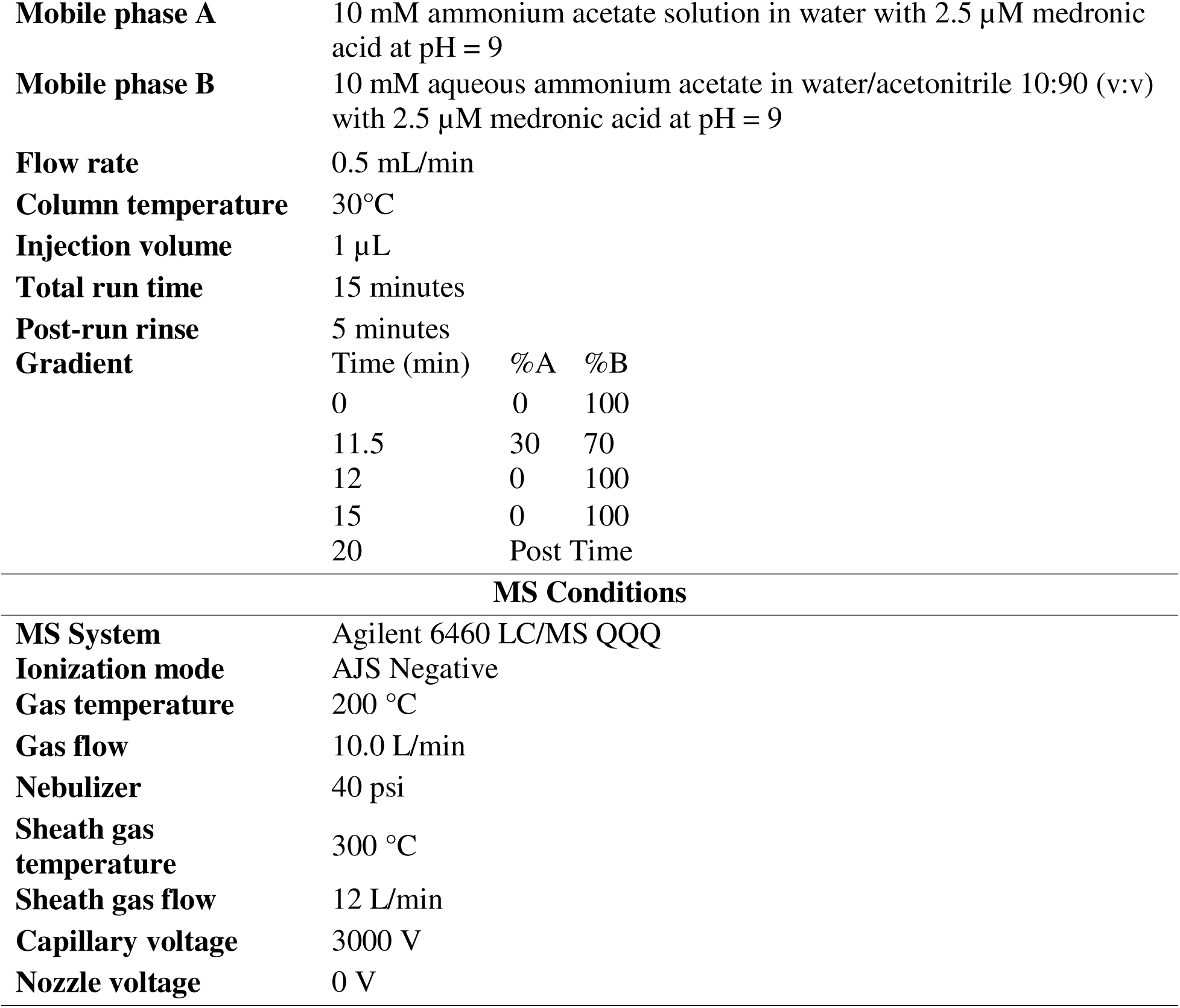
Table 1.

**Table 1.**
| HPLC Conditions |  |
| --- | --- |
| **Column** | Agilent InfinityLab Poroshell 120 HILIC-Z, PEEK lined, 2.1 × 100 mm, 2.7 µm, (p/n 685775-924) |

|  |  |  |  |
| --- | --- | --- | --- |
| Mobile phase A | 10 mM ammonium acetate solution in water with 2.5 μM medronic acid at pH = 9 |  |  |
| Mobile phase B | 10 mM aqueous ammonium acetate in water/acetonitrile 10:90 (v:v) with 2.5 μM medronic acid at pH = 9 |  |  |
| Flow rate | 0.5 mL/min |  |  |
| Column temperature | 30°C |  |  |
| Injection volume | 1 μL |  |  |
| Total run time | 15 minutes |  |  |
| Post-run rinse | 5 minutes |  |  |
| Gradient | Time (min) | %A | %B |
|  | 0 | 0 | 100 |
|  | 11.5 | 30 | 70 |
|  | 12 | 0 | 100 |
|  | 15 | 0 | 100 |
|  | 20 | Post Time |  |
| MS Conditions |  |  |  |
| MS System | Agilent 6460 LC/MS QQQ |  |  |
| Ionization mode | AJS Negative |  |  |
| Gas temperature | 200 °C |  |  |
| Gas flow | 10.0 L/min |  |  |
| Nebulizer | 40 psi |  |  |
| Sheath gas temperature | 300 °C |  |  |
| Sheath gas flow | 12 L/min |  |  |
| Capillary voltage | 3000 V |  |  |
| Nozzle voltage | 0 V |  |  |

**Table 2.** Instrument Conditions.

| **Molecule** | **Retention time (min)** | **Precursor ion (m/z)** | **Product ion (m/z)** | **Product ion Collision energy (V)** | **Fragmentor (V)** |
| --- | --- | --- | --- | --- | --- |
| Itaconic Acid | 7.87 | 129 | 85 | 8 | 115 |
|  | 7.87 | 129 | 41 | 12 | 115 |
| Citraconic Acid | 1.69 | 129 | 85 | 8 | 115 |
|  | 1.69 | 129 | 41 | 12 | 115 |
| Mesaconic Acid | 9.47 | 129 | 85 | 8 | 115 |
|  | 9.47 | 129 | 41 | 12 | 115 |

The acquired raw files were analyzed using Agilent MassHunter Workstation Software (Agilent Technologies, CA, USA) according to Agilent familiarization guides (Qualitative analysis G3335-90198; Quantitative analysis G3335-90152). In the Qualitative Analysis Navigator (Version B.08.00) program, the pressure profiles were extracted, and chromatograms were checked to ensure no significant pressure and retention time shift. Then, specific retention time for itaconic acid, citraconic acid and mesaconic acid was obtained in the Qualitative Analysis Workflow (Version B.08.00) program. The quantitative analysis was conducted in Agilent MassHunter Quantitative Analysis for QQQ (Version B.08.00/Build 8.0.598.0), using information including retention time, paired precursor and product ions, and standard concentrations. After the method application, calibration curves were plotted using standard levels, and the concentration of target molecules in samples was determined. The separation of itaconic acid, citraconic acid, and mesaconic acid during the sample analysis in the standard (five standard levels are shown as an example) can be seen in Figure 1.

**Table 3.** Antibodies Utilized.

| **Antibody** | **Colour** | **Clone** |
| --- | --- | --- |
| CD4 | PE | GK 1.5 |
| CD8 | FITC | 53-6.7 |
| CD44 | BV480 | IM7 |
| CD25 | PE-TexasRed | PC61(RUO) |
| RORγt | APC | B2D |
| Tbet | QDot 800 | 4B10 |
| FOXP3 | Pacific Blue | FJK-165 |
| GATA3 | BV711 | L50-823 |

### Statistical Analysis

Data are expressed as average ± standard deviation (avg±s.d.) or average ± standard error (avg±s.e.). Comparisons between multiple treatment groups were performed using one-way ANOVA with Fisher’s LSD test and P-values of smaller than 0.05 were considered statistically significant. Comparisons between two treatment groups were performed by two-tailed T test and P-values of smaller than 0.05 were considered statistically significant (GraphPad Prism Software 10.0, San Diego, CA).

## Acknowledgement

The author acknowledge support from Northeast Ohio High Field NMR facility, Chemical Engineering Instrumentation Core for NMR and FTIR support. We acknowledge the Cleveland Clinic Research Proteomics and Metabolomics Core Facility (RRID:SCR_026563), especially Dr Belinda Willard, for their assistance with global proteomics. We also thank the support from Cleveland DDRCC Histology core for all histology imaging, especially from Dr. Carlo De Salvo. The authors acknowledge funding from Coulter Foundation at Case Western Reserve University.

## Author Contribution

HQ designed the study, performed experiments, and wrote the manuscript. DS, HQ, ZC and WZ performed *in vivo* experiment. AA, HQ, TK and ZC performed data analysis. HQ, ZC, WZ, AT, MMCSJ, SP, APS, PM, JS, VR, and TK performed *ex vivo* experiments. CDS and DS performed histology. HQ and BZ performed LC-MS/MS. YA provided plasmid. AA and AB provide the idea and guiding.

